# Advancements in Cardiovascular Disease Detection: Leveraging Data Mining and Machine Learning

**DOI:** 10.1101/2024.03.09.584222

**Authors:** Md. Sahadat Hossain, Md. Alamin Talukder, Md. Zulfiker Mahmud

## Abstract

Cardiovascular disease (CVD) is a significant global health concern, requiring early detection and accurate prediction for effective intervention. Machine learning (ML) offers a data-driven approach to analyzing patient data, identifying complex patterns and predicting CVD risk factors like blood pressure (BP), cholesterol levels, and genetic predispositions. Our research aims to predict CVD presence using ML algorithms, leveraging the Heart Disease UCI dataset with 14 attributes and 303 instances. Extensive feature engineering enhanced model performance. We developed five models using Logistic Regression, K-Nearest Neighbors (KNN), Decision Tree Classifier, Support Vector Machine (SVM), and Random Forest Classifier, refining them with hyperparameter tuning. Results show substantial accuracy improvements post-tuning and feature engineering. ‘Logistic Regression’ achieved the highest accuracy at 93.44%, closely followed by ‘Support Vector Machine’ at 91.80%. Our findings emphasize the potential of ML in early CVD prediction, underlining its value in healthcare and proactive risk management. ML’s utilization for CVD risk assessment promises personalized healthcare, benefiting both patients and healthcare providers. This research showcases the practicality and effectiveness of ML-based CVD risk assessment, enabling early intervention, improving patient outcomes, and optimizing healthcare resource allocation.

## 1. Introduction

Cardiovascular Disease (CVD) remains one of the most pervasive and lethal health challenges worldwide, claiming the lives of 17.9 million individuals annually, representing 31% of all global deaths [9]. This grim statistic underscores the pressing need for innovative approaches to combat this disease.

The advent of data mining as a burgeoning field has ushered in a new era for uncovering concealed patterns within vast datasets [14, 21]. In parallel, the clinical sciences generate an abundance of data, including clinical reports and a myriad of patient symptoms. Leveraging data mining and machine learning techniques, we can address critical prediction-related challenges within the realm of clinical fields, particularly those associated with cardiovascular health [15].

Machine learning stands as the linchpin of this transformative process, enabling the extraction of latent patterns from clinical data and facilitating predictions for the future [8]. These concealed insights gleaned from clinical datasets hold great promise for advancing medical diagnostics and care. Yet, a formidable obstacle remains—clinical datasets are characterized by their widespread distribution, inherent heterogeneity, and colossal scale. Bridging this divide calls for their seamless integration into hospital management systems [18, 19].

Within the realm of prediction-oriented challenges, a pantheon of data mining and machine learning algorithms and techniques awaits exploration [20, 22, 17, 16, 5, 10]. In this study, we harness the power of five distinct machine-learning algorithms and data mining techniques. Our overarching aim is to develop predictive models that efficiently detect cardiovascular disease. This strategic pursuit is motivated by a simple yet profound imperative—saving lives by enabling quicker and more accurate treatment interventions [3].

The term ‘cardiovascular disease’ encompasses a spectrum of heart-related conditions, such as heart attacks, angina, and strokes, contributing to a significant proportion of global mortality. Timely detection of these insidious conditions assumes paramount importance in averting potentially fatal outcomes. In our contemporary digital era, healthcare institutions, including medical centers and hospitals, generate vast volumes of data daily. It is within this deluge of data that machine learning and data mining algorithms and techniques hold the potential to discern patterns and predict the occurrence of cardiovascular disease in patients.

While existing research efforts have addressed the challenge of cardiovascular disease prediction, we embark on a distinctive journey. To our knowledge, no previous study has ventured to construct and compare five diverse prediction models for cardiovascular disease. This research employs five unique machine learning algorithms, subjecting each to rigorous training and testing on validated datasets. The outcome of this endeavor is a comprehensive comparison of algorithmic performance, illuminating which among them proves most effective. Through meticulous feature engineering and hyperparameter tuning, we aim to push the boundaries of accuracy in cardiovascular disease prediction [1].

The methods applied in this research encompass Logistic Regression, K-Nearest Neighbors (KNN), Support Vector Machine (SVM), Decision Tree Classifier, and Random Forest Classifier.

At the core of this research lies a dual objective. First and foremost, we aspire to construct machine learning models capable of predicting the presence of cardiovascular disease in patients with unparalleled accuracy. Simultaneously, we strive to optimize the predictive process, curbing the number of tests required and the time expended in making diagnoses. Our specific objectives are delineated as follows:

- Create Cardiovascular Disease prediction models by leveraging a spectrum of Machine Learning algorithms and Data Mining techniques, including Support Vector Machine, Decision Tree Classifier, Logistic Regression, Random Forest Classifier, and K-Nearest Neighbor.
- Predict the presence of cardiovascular disease in patients by deploying diverse machine learning algorithms on certified datasets.
- Identify correlations between a myriad of patient characteristics and attributes to enhance predictive precision.
- Elevate the standard of cardiovascular disease prediction, thereby diminishing the quantity of tests required and the time necessary for diagnosis through comprehensive analysis and comparison of results from various machine learning models.

The forthcoming sections of this paper are meticulously organized. Section 2 provides a concise survey of related literature, contextualizing our research within the existing body of knowledge. Section 3 presents the methodology we have devised and offers insights into the datasets we employ. Section 4 delves into the experimental setup and furnishes a comprehensive evaluation of our deep learning model. Lastly, in Section 9, we draw our conclusions and outline avenues for future research within this vital domain.

## 2. Related works

In machine learning, various algorithms serve as indispensable tools for predictive modeling across various domains. This research is dedicated to the development of models designed to predict the presence of cardiovascular heart disease in patients. To accomplish this, we have crafted five distinct machine learning models, each harnessing a different algorithmic approach, all of which have been applied to the UCI dataset[7].

The rationale behind employing five diverse machine learning algorithms is to enable a comprehensive evaluation of their effectiveness in addressing this specific problem. Such an approach not only facilitates the identification of the most suitable algorithm for this task but also provides a basis for comparing and contrasting the performance of these models. This strategy ensures that our findings are robust and grounded in thoroughly exploring the available algorithmic options. The five machine learning algorithms adopted for this research include: 1. Decision Tree 2. Logistic Regression 3. Support Vector Machine (SVM) 4. Random Forest 5. K-Nearest Neighbor The selection of these algorithms is informed by the need to explore various computational approaches to the problem at hand, ensuring a comprehensive and methodical investigation. It is noteworthy that prior research has explored the application of machine learning algorithms for heart disease or cardiovascular disease prediction. In the following, we delve into these earlier studies, highlighting their insights and contributions, which inform and complement the research presented here.

B. VenkataLakshmi et al. [6] designed and developed a heart disease prediction system using machine learning algorithms. They used the Cleveland heart disease dataset from the UCI Machine Learning Repository, which contains 13 attributes and 294 patient records. Two classification algorithms were implemented - Decision Tree Classifier and Naive Bayes. The performance of these models was evaluated and Naive Bayes achieved a slightly higher accuracy of 85.034% compared to 84.013% for Decision Tree. They proposed applying genetic algorithm for attribute selection as future work to potentially improve performance.

Jaymin Patel et al. [13] implemented three popular machine learning algorithms - J48 decision tree, Logistic Model Tree and Random Forest on the Cleveland heart disease dataset in 2015. 10-fold cross validation was used to evaluate the predictive performance of these models. Among the classifiers, J48 with reduced error pruning achieved the highest test accuracy of 56.76%.

K. Gomath et al. [11] developed machine learning models to predict heart disease in male patients. They used Naive Bayes, J48 decision tree and Artificial Neural Network (ANN) classifiers on a dataset containing 8 attributes and records of 210 male patients. Naive Bayes yielded the best results with an accuracy of 79.90%.

Zeinab Arabasadi et al. [2] proposed a hybrid diagnosis model for coronary artery disease in 2017, combining Artificial Neural Network and genetic algorithm. They applied their methodology on a dataset of 303 patients with 22 relevant clinical attributes. The combined ANN-Genetic algorithm model achieved the highest predictive accuracy of 93.85

Sarangam Kodati et. al [12] implemented Random Forest, Decision Table, Naive Bayes and J48 classification algorithms on a heart disease dataset comprising 303 records and 14 attributes in 2018. Their results showed that Decision Table performed best with an accuracy of 84.81%.

Bharti. Rohit et al. [4] employed diverse machine learning algorithms and applied deep learning to compare the results and analyze the UCI Machine Learning Heart Disease dataset. The dataset comprised 14 key attributes utilized for the analysis. Substantial and promising results were achieved, and these findings were validated using accuracy metrics and a confusion matrix. To enhance the analysis, they addressed irrelevant features in the dataset through the implementation of Isolation Forest. Additionally, data normalization was performed to optimize results. The study explored the integration of this analysis with multimedia technology, particularly on mobile devices. Through the application of a deep learning approach, an accuracy of 94.2% was obtained.

To summarize, various machine learning algorithms have been applied for cardiovascular disease prediction on standard datasets. Ensemble and hybrid models have achieved better performance compared to individual classifiers. Attribute selection and optimization techniques can be explored to improve the prediction accuracy further.

## 3. Methodology

The methodology adopted for this study involved applying machine learning algorithms to predict heart disease using clinical data. A systematic approach was followed which can be summarized in the block diagram shown in Figure 1.

**Figure 1:**
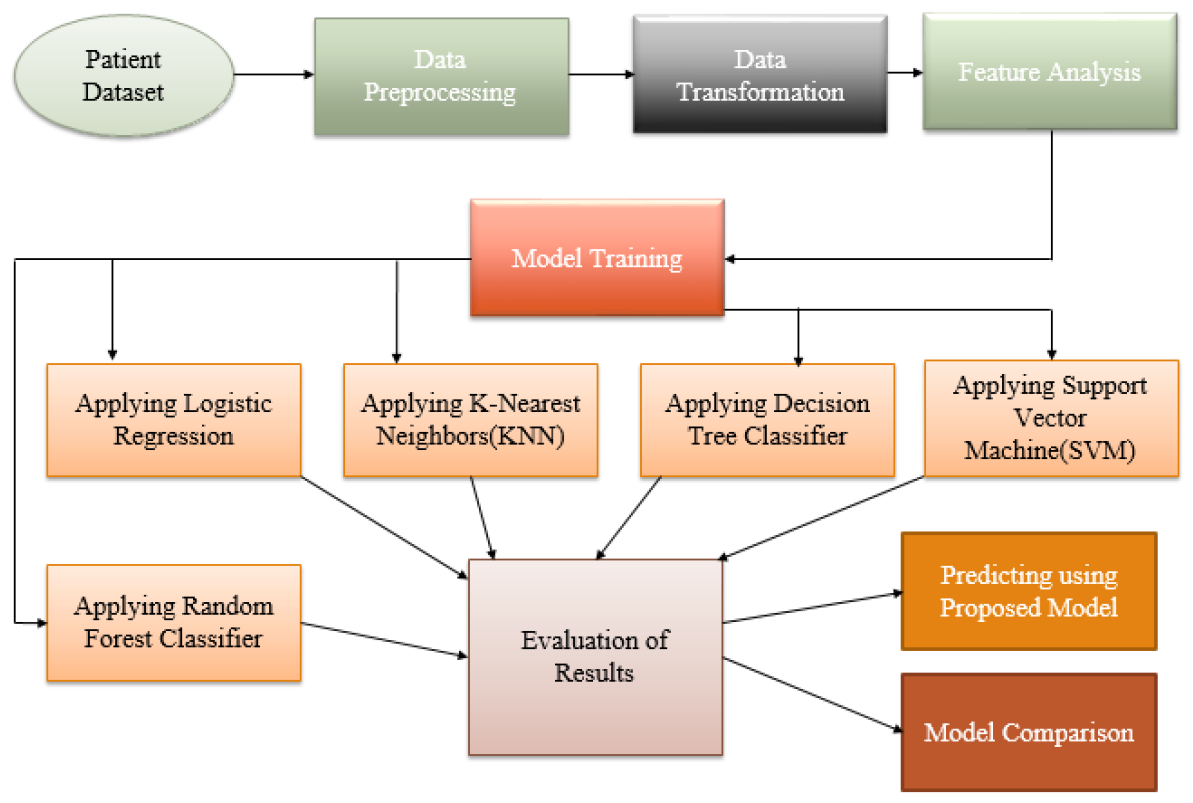
Methodology workflow adopted for the study

The main steps were:

1. **Data Collection**: Relevant heart disease datasets were obtained from online repositories like UCI Machine Learning Repository.
2. **Data Preprocessing**: The datasets were examined for completeness, consistency and outliers. Missing or incorrect values were addressed. Features were encoded and normalized.
3. **Model Building**: Supervised machine learning algorithms like Decision Trees, Naive Bayes, Neural Networks etc. were implemented using the preprocessed datasets.
4. **Model Evaluation**: Models were evaluated using performance metrics like accuracy, sensitivity, specificity obtained through validation techniques like k-fold cross validation.
5. **Model Comparison**: Top performing models were identified and their predictive ability was compared. Parameter tuning was also performed.
6. **Conclusion**: The best model for heart disease prediction was selected based on evaluation. Opportunities for improvement and future work were identified.

The above workflow formed the methodological framework of this study. Relevant details of data, algorithms and evaluation metrics used are provided in subsequent sections.

### 3.1. Dataset

The heart disease dataset from the UCI Machine Learning Repository [7] was used in this study. It is a widely used benchmark dataset for evaluating heart disease prediction models. The dataset contains 303 records with 14 clinically measured attributes from patients. The attributes include demographic details (age, sex), medical test results (e.g. cholesterol, blood pressure), symptoms (e.g. chest pain type), and the presence/absence of heart disease (target variable). A description of each attribute is provided in Table 1. The target variable indicates if heart disease is present (value 1) or not (value 0). This variable was used to develop supervised learning models. The dataset was randomly split into 70% for the training set and 30% for the testing set. The training set was used to build the machine learning models while the testing set served to evaluate the predictive performance of these models. This UCI heart disease dataset is widely employed in related research work due to its benchmark status, size and variety of features that capture different dimensions of cardiac health status. The availability of a clearly defined target variable also makes it suitable for supervised learning approaches.

**Table 1:** The brief description of the dataset attributes.

| Attributes | Meaning and Values |
| --- | --- |
| age | Age in years |
| sex | (1 = male, 0 = female) |
| cp | chest pain type (4 values) |
| trestbps | resting blood pressure (in mm Hg) |
| chol | serum cholestoral (in mg/dl) |
| fbs | fasting blood sugar > 120 mg/dl (1 = true; 0 = false) |
| restecg | resting electrocardiographic results (values 0,1,2) |
| thalach | maximum heart rate achieved |
| exang | exercise induced angina |
| oldpeak | ST depression induced by exercise relative to rest |
| slope | the slope of the peak exercise ST segment |
| ca | number of major vessels (0-3) colored by flourosopy |
| thal | 4 types of values (0, 1, 2, 3) |
| target | (0 = no presence, 1 = presence) |

### 3.2. Data Preprocessing

After collecting the dataset, the first task is to preprocess the data before using it. Data Preprocessing is a data processing technique that involves several steps to complete the process, including changing the data format and some other steps. In general, real-world data is often inconsistent, incomplete, and missing in certain behaviors, and contains many errors and outliers. By data preprocessing, we can solve all of these problems and make the dataset ready for use in any machine learning model. When we train a machine learning model with a dataset that is not preprocessed, the model will not be able to give better results, and the performance of that model will eventually decrease significantly. Here are the steps for data preprocessing that we have followed in this research:

i. Identifying and removing duplicate rows is the first step of data preprocessing.
ii. Rows containing missing values were subsequently identified and removed.
iii. After that, box and whisker plots were used to find the outliers, and
iv. Subsequently, remove those rows with outliers.
v. Find out and fill in the missing values in the dataset.

In Figure 2, we present the initial data frame of the dataset. This visual representation reveals that the dataset consists of 14 attributes or columns and 303 rows or instances. Notably, all attribute values are already in numerical format, eliminating the need for any conversion.

**Figure 2:**
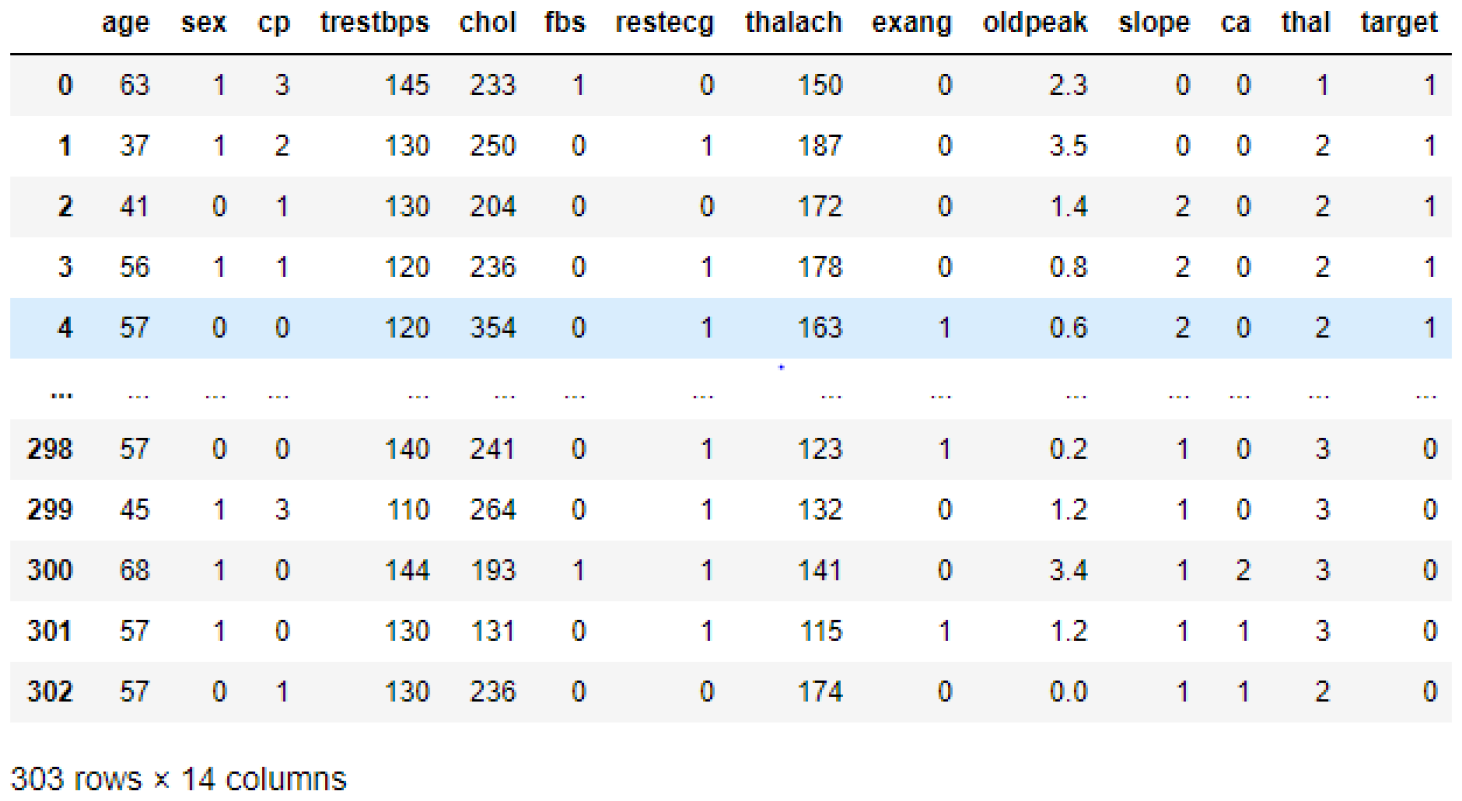
Initial Data frame of the Dataset

### 3.3. Feature Analysis

Feature analysis is one of the most important thing in any machine learning related problem solving. Which feature is more important, which feature is less, how are the features interrelated to the target value and many more things can be cleared by doing feature analysis. In this section we are going to discuss about the feature analysis, how we choose the best features and other things clearly. After reading the data set in csv file the first thing that we did is data preprocessing. In data preprocessing we have checked whether there is any missing data, outliers, or inconsistent data in the data set. After doing data preprocessing we analysis the features. We have seen that there were no missing value in the data set and it seemed the dataset were pretty balanced as there were 165 records whose target value 1 and 138 records whose value 0. We have separated the categorical and continuous value. We have plotted the correlation matrix and have seen how the features or attributes are interrelated with each other. We also have plotted several diagram for every feature and did analysis for each and every thing for getting better picture of the features relation in the data set.

### 3.4. Feature Engineering

In the data set there can be many features that can affect the accuracy of the algorithm in positive or negative way. That is why working with the feature is very important. There can be situation that we need to work with some feature and we need to reduce some other feature. If we can reduce some feature it will help us to train our models faster. We can even make new feature by dividing or interacting with existing features, sometimes this give huge amount of improvement in our models. Feature Engineering means working with the features, modifying them and choosing the best features for the model training. Using the domain knowledge to extract features from the raw data via data mining techniques and machine learning knowledge is feature engineering. To improve the accuracy and performance of the machine learning algorithms these feature can be used. we have splitted the features whose value is categorical and added dummy features, because we have seen doing this we got better performance.

### 3.5. Feature Importance

After doing feature analysis and feature engineering the next question that comes in our mind is that what features have the biggest impact on the target value or on the predictions. Finding the importance of the features is called feature importance. In the data set there can have some features or attributes which do not have much impact on the prediction, on the other hand in some cases there can be some attributes which can reduce the models accuracy and performance. That is why it is very important to find out the important features and working with those features for training the models. One thing is need to remember that feature importance can be different for different machine learning algorithms. We have checked the feature importance before training my models. We used Univariate feature selection method for getting the feature importance. It selects the best features based on univariate statistical tests. We did it using SciKit learn feature selection function. By this we have seen the importance value of every attribute or feature.

### 3.6. Feature Selection

After getting the feature importance then the task that need to done is feature selection. There can be some features which are related to others in linearly. This can put pressure on the models and reduce the models performance. Feature selection refers to select the important features that are related to the model performance and reducing unnecessary features that can harm model performance. Unnecessary features reduce the training and testing time and also reduce the models performance and accuracy. We can reduce the model training and testing time and also reduce the over-fitting problem by doing feature selection. After doing univariate feature selection method and select-k-based algorithm, we have chosen the best features for training my models. We have removed four lowest important features.

### 3.7. Performance Metrics

A Confusion Matrix, also known as an error matrix, provides a visual representation of the performance of a machine learning algorithm, especially in supervised learning. It is a two-dimensional contingency table, where the actual and predicted class represent the x and y axes, respectively.

In Figure 3, a depiction of the confusion matrix is presented. The matrix is composed of four components: True Positives (TP), True Negatives (TN), False Positives (FP), and False Negatives (FN).

**Figure 3:**
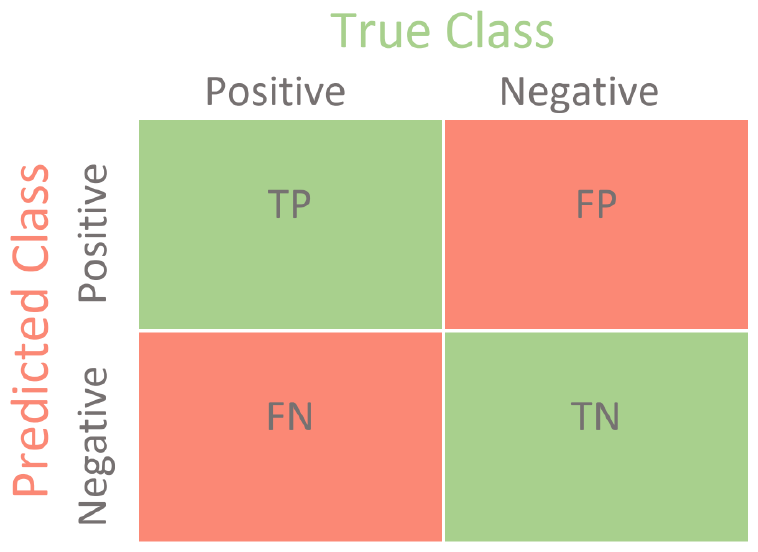
Confusion Matrix

The confusion matrix is instrumental in calculating various performance metrics, including accuracy, precision, sensitivity, and specificity. The formulas for these metrics are provided below:

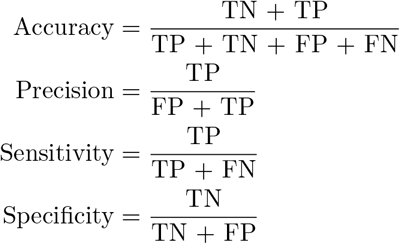

### 3.8. Hyper Parameter Tuning

In machine learning, we can use various different algorithms. In every algorithm there are some parameters that the algorithm takes and works according to the parameters. The performance of the algorithms can vary because of the parameters values. The hyper parameter tuning means that choosing the values of the parameters of any machine learning algorithm by which that algorithm gives better accuracy and result. The value of the parameters actually controls the performance of the algorithm and the training-testing process.

Hyper parameter tuning is very important in the machine learning model training and testing, because it controls the performance of the algorithms. That is why experts put a lot of effort for finding the best combination of the hyper parameters. There are various way for hyper parameter tuning, we can follow those way for doing hyper parameter tuning. The most common and used hyper parameter tuning methods are Random search, Grid search etc. There are many others but they also take a lot of effort and these two are the most used methods. We have applied these two methods in this research for doing hyper parameter tuning.

### 3.9. Modeling and Predicting

The main purpose of this research is to predict cardiovascular disease or heart disease with highest accuracy. For doing this we have made five different models using five different and most used machine learning algorithms. In the chapter 4 we will discuss about the results of my models and compare my models in terms of their accuracy and performance. Here in this section we will discuss about the model building and training and testing related tasks. To build, train and test my models the tools that helped me most is SciKit Learn library. we did the coding part in python programming language. We have splitted the data set in two part, one is training data set and another is testing data set. The ratio was 80:20, 80% of the full data set is used for training purpose and 20% is used for testing purpose. Then we created the models using built in functions and run all five different algorithms.

### 3.10. Finding the result

After creating the models, we train the models with the training data set then we tested the models with the testing data set and got the results for every models. Next chapter we are going to show the result part very clearly and details with creating the summary table. We run the models basically in two steps, first we trained and test the models with no hyper parameter tuning, secondly we did hyper parameter tuning for the algorithms and then trained and test. After hyper parameter tuning the performance of the models improved significantly. Simpler models or methods proved to be very useful by giving decent performance. We have given details explanation about the models performance in the later chapter.

## 4. Result Analysis

The results presented in this section stem from experiments employing various machine learning algorithms to predict the presence of cardiovascular disease. Two experiments were conducted: the first without hyperparameter tuning, and the second after hyperparameter tuning.

### 4.1. Without Hyperparameter Tuning

#### 4.1.1. Models on Training

After data preprocessing and feature engineering, models were trained and tested without hyperparameter tuning. Table 2 provides the precision, recall, and F1 scores of various machine learning models during training without hyperparameter tuning. Notably, the Logistic Regression model exhibits high precision, recall, and F1 scores, indicating a balanced performance in correctly identifying positive and negative instances. However, all models achieved perfect scores in this scenario, suggesting potential overfitting.

**Table 2:**
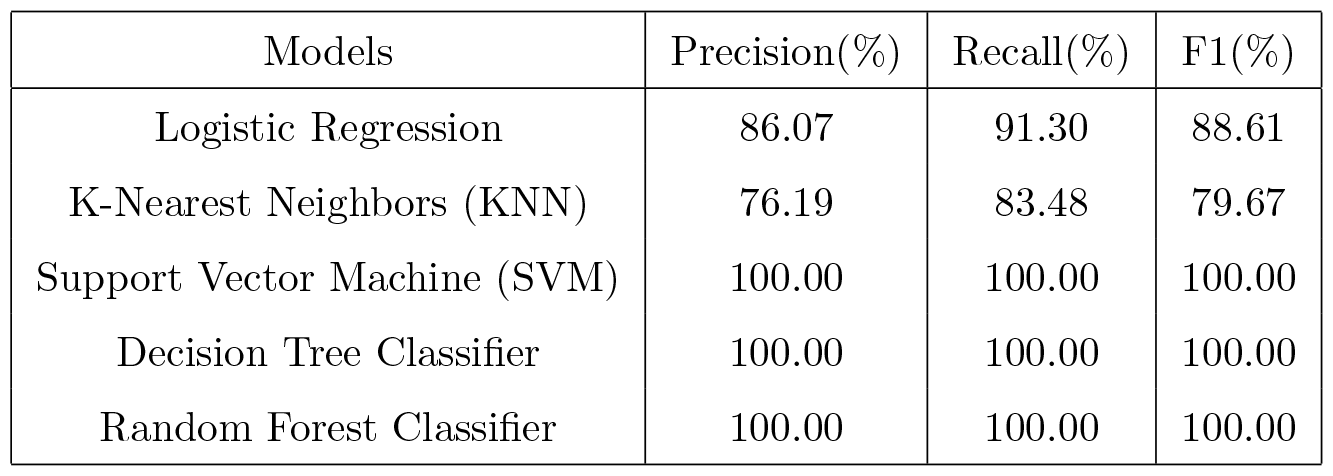
Classification report of models during training without hyperparameter tuning.

| Models | Precision(%) | Recall(%) | F1(%) |
| --- | --- | --- | --- |
| Logistic Regression | 86.07 | 91.30 | 88.61 |
| K-Nearest Neighbors (KNN) | 76.19 | 83.48 | 79.67 |
| Support Vector Machine (SVM) | 100.00 | 100.00 | 100.00 |
| Decision Tree Classifier | 100.00 | 100.00 | 100.00 |
| Random Forest Classifier | 100.00 | 100.00 | 100.00 |

Table 3 showcases the accuracy achieved during training without hyperparameter tuning. Training accuracy highlights the overall performance of each model during the training phase. While all models achieved high accuracy, the Support Vector Machine (SVM), Decision Tree Classifier, and Random Forest Classifier show perfect accuracy. It is essential to consider potential overfitting due to the large discrepancy between training and testing accuracy.

**Table 3:**
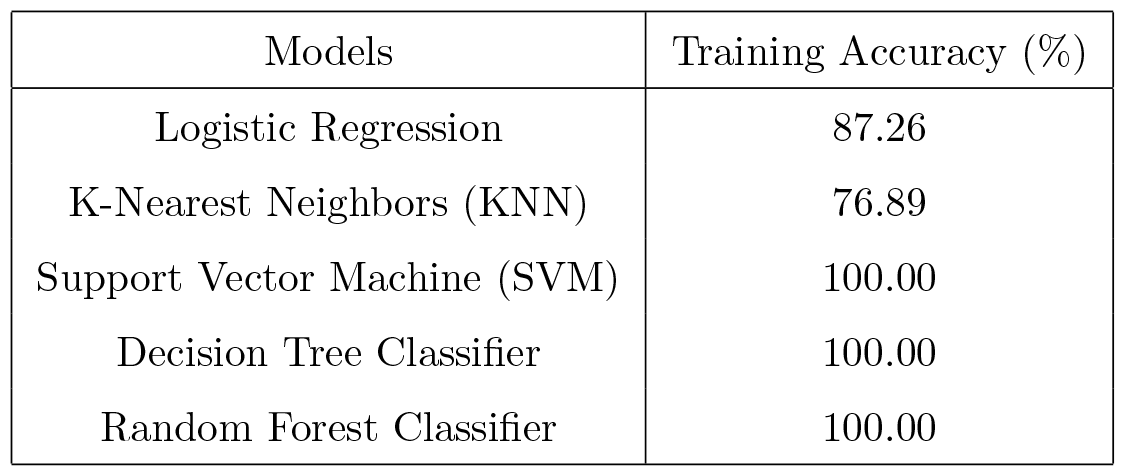
Models performance on training without hyperparameter tuning.

Table 4 illustrates the confusion matrix during training without hyperparameter tuning. The confusion matrix provides a detailed breakdown of true positives (TP), false positives (FP), false negatives (FN), and true negatives (TN) for each model during training. The SVM, Decision Tree, and Random Forest classifiers exhibit no false positives or false negatives, indicating robust performance in classifying instances.

**Table 4:** Confusion Matrix of training without hyperparameter tuning.

| Models | TP | FP | FN | TN |
| --- | --- | --- | --- | --- |
| Logistic Regression | 78 | 19 | 8 | 107 |
| K-Nearest Neighbors (KNN) | 67 | 30 | 19 | 96 |
| Support Vector Machine (SVM) | 97 | 0 | 0 | 115 |
| Decision Tree Classifier | 97 | 0 | 0 | 115 |
| Random Forest Classifier | 97 | 0 | 0 | 115 |

#### 4.1.2. Models on Testing

Table 5 provides the classification report during testing without hyperparameter tuning. In the testing phase without hyperparameter tuning, the classification report shows varied performance across models. Logistic Regression demonstrates a balanced performance with good precision, recall, and F1 scores, while Support Vector Machine exhibits high recall but lower precision. K-Nearest Neighbors shows lower overall performance.

**Table 5:** Classification report of models on testing without hyperparameter tuning.

| Models | Precision(%) | Recall(%) | F1(%) |
| --- | --- | --- | --- |
| Logistic Regression | 82.35 | 84.00 | 83.17 |
| K-Nearest Neighbors (KNN) | 67.92 | 72.00 | 69.90 |
| Support Vector Machine (SVM) | 54.95 | 100.00 | 70.92 |
| Decision Tree Classifier | 79.55 | 70.00 | 74.47 |
| Random Forest Classifier | 82.35 | 84.00 | 83.17 |

Table 6 showcases the accuracy achieved during testing without hyperparameter tuning. Testing accuracy provides insights into the generalization performance of models. Logistic Regression and Random Forest Classifier outperform other models in testing accuracy, but the significant drop from training accuracy suggests potential overfitting issues.

**Table 6:** Models performance on testing without hyperparameter tuning.

| Models | Testing Accuracy (%) |
| --- | --- |
| Logistic Regression | 81.32 |
| K-Nearest Neighbors (KNN) | 65.93 |
| Support Vector Machine (SVM) | 54.95 |
| Decision Tree Classifier | 73.63 |
| Random Forest Classifier | 81.32 |

Table 7 illustrates the confusion matrix during testing without hyperparameter tuning. The confusion matrix provides a detailed breakdown of true positives (TP), false positives (FP), false negatives (FN), and true negatives (TN) for each model during training. The SVM, classifier exhibits no false positives or false negatives, indicating robust performance in classifying instances.

**Table 7:** Confusion Matrix of testing without hyperparameter tuning.

| Models | TP | FP | FN | TN |
| --- | --- | --- | --- | --- |
| Logistic Regression | 32 | 9 | 8 | 42 |
| K-Nearest Neighbors (KNN) | 24 | 17 | 14 | 36 |
| Support Vector Machine (SVM) | 0 | 41 | 0 | 50 |
| Decision Tree Classifier | 31 | 10 | 14 | 36 |
| Random Forest Classifier | 32 | 9 | 8 | 42 |

### 4.2. With Hyperparameter Tuning

Hyperparameter tuning is crucial for model generalization and avoiding overfitting. In this section, we discuss the results after applying hyperparameter tuning.

#### 4.2.1. Models on Training

Table 8 provides the classification report during training with hyperparameter tuning. After hyperparameter tuning, the models exhibit improved precision, recall, and F1 scores during training. Notably, the SVM, Decision Tree, and Random Forest models continue to demonstrate excellent performance.

**Table 8:** Classification report of models on training with hyperparameter tuning.

| Models | Precision(%) | Recall(%) | F1(%) |
| --- | --- | --- | --- |
| Logistic Regression | 86.53 | 91.73 | 89.05 |
| K-Nearest Neighbors (KNN) | 78.19 | 85.48 | 83.67 |
| Support Vector Machine (SVM) | 64.09 | 87.22 | 73.89 |
| Decision Tree Classifier | 100.00 | 100.00 | 100.00 |
| Random Forest Classifier | 100.00 | 100.00 | 100.00 |

Table 9 showcases the accuracy achieved during training with hyperparameter tuning. Training accuracy remains high after hyperparameter tuning, with Logistic Regression and Random Forest Classifier achieving the highest accuracy. The improvements suggest better model convergence.

**Table 9:** Models performance on training with hyperparameter tuning.

| Models | Training Accuracy (%) |
| --- | --- |
| Logistic Regression | 87.78 |
| K-Nearest Neighbors (KNN) | 82.64 |
| Support Vector Machine (SVM) | 85.54 |
| Decision Tree Classifier | 83.88 |
| Random Forest Classifier | 91.32 |

Table 10 illustrates the confusion matrix during training with hyperparameter tuning. In the confusion matrix, models demonstrate improved true positive rates and reduced false positive rates after hyperparameter tuning. The Random Forest Classifier stands out with fewer false positives.

**Table 10:** Confusion Matrix of training with hyperparameter tuning.

| Models | TP | FP | FN | TN |
| --- | --- | --- | --- | --- |
| Logistic Regression | 79 | 18 | 8 | 107 |
| K-Nearest Neighbors (KNN) | 76 | 18 | 19 | 99 |
| Support Vector Machine (SVM) | 75 | 18 | 12 | 107 |
| Decision Tree Classifier | 79 | 19 | 15 | 99 |
| Random Forest Classifier | 79 | 9 | 9 | 115 |

#### 4.2.2. Models on Testing

Table 11 provides the classification report during testing with hyperparameter tuning. Hyperparameter tuning enhances the classification metrics during testing. Logistic Regression and Decision Tree Classifier show balanced precision, recall, and F1 scores, addressing the overfitting observed in the untuned models.

**Table 11:** Classification report of models on testing with hyperparameter tuning.

| Models | Precision(%) | Recall(%) | F1(%) |
| --- | --- | --- | --- |
| Logistic Regression | 90.62 | 90.62 | 90.62 |
| K-Nearest Neighbors (KNN) | 68.57 | 75.00 | 71.64 |
| Support Vector Machine (SVM) | 66.67 | 87.50 | 75.68 |
| Decision Tree Classifier | 83.87 | 81.25 | 82.54 |
| Random Forest Classifier | 87.50 | 87.50 | 87.50 |

Table 12 showcases the accuracy achieved during testing with hyperparameter tuning. Testing accuracy after hyperparameter tuning reflects the models’ improved ability to generalize to unseen data. Logistic Regression and SVM emerge as top performers in accuracy, indicating effective hyperparameter adjustments.

**Table 12:**
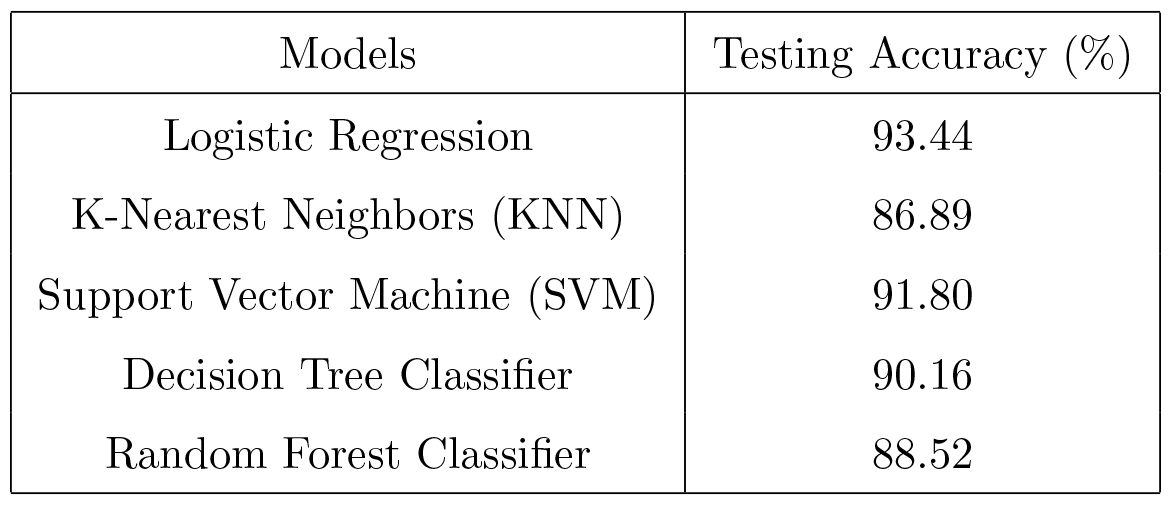
Models performance on testing with hyperparameter tuning.

| Models | Testing Accuracy (%) |
| --- | --- |
| Logistic Regression | 93.44 |
| K-Nearest Neighbors (KNN) | 86.89 |
| Support Vector Machine (SVM) | 91.80 |
| Decision Tree Classifier | 90.16 |
| Random Forest Classifier | 88.52 |

Table 13 illustrates the confusion matrix during testing with hyperparameter tuning. In the confusion matrix, models demonstrate improved true positive rates and reduced false positive rates after hyperparameter tuning. The Random Forest Classifier stands out with fewer false positives.

**Table 13:** Confusion Matrix of testing with hyperparameter tuning.

| Models | TP | FP | FN | TN |
| --- | --- | --- | --- | --- |
| Logistic Regression | 83 | 7 | 7 | 115 |
| K-Nearest Neighbors (KNN) | 77 | 18 | 10 | 107 |
| Support Vector Machine (SVM) | 80 | 8 | 9 | 115 |
| Decision Tree Classifier | 77 | 9 | 11 | 115 |
| Random Forest Classifier | 73 | 15 | 9 | 115 |

## 5. Cross Validation

In machine learning model training, Cross validation is a very important step. By doing this we can track our model so that we can know about the risk of over-fitting. We can trust our CV(Cross validation) scores. Normally 5-fold CV is enough for machine learning models. The CV scores will be more reliable if we use more folds. But If we increase the folds the training takes longer time to finish. When the training data is limited we should not use too many folds. If we use too many folds we would have too few sample in each fold to guarantee statistical significance. Often the data is imbalanced, i.e. there may be less number of class instances but high number of class1 instances. Therefore we need to train and test our algorithms on every instances of the dataset. After that we can take an average of all the noted accuracies over the data set. The K-fold cross validation first divides the data set into k-subsets. Suppose, we have divided the dataset into k=10 subsets. It reserve 1 part for testing the algorithms or models and rest of the parts for train the algorithms. It continue the process by changing the testing part in each iteration and train the algorithm over the other parts. Then the errors and accuracies are averaged to get an average accuracy and errors of the algorithms. An algorithm can be under-fit or over-fit over a data set for some training data and testing data set. But If we do cross validation we can achieve a generalized model. Therefore we have used K-fold cross validation in our research.

## 6. AUC-ROC Curve

The measurement of machine learning models performance is very important task in machine learning researches. For this performance measurement the most used way on which we can rely is AUC-ROC curve. This is mainly used in classification related problems. The AUC (Area Under the Curve) and ROC (Receiver Operating Characteristics) curve is used in multi class classification problem. There are two things combined in one diagram, first one is ROC which is a probability curve and the measure of separability is represented by AUC. The higher the AUC the higher the model performance. Basically it means that the model can predict more accurately. This curve is plotted where on the x-axis the FPR (False Positive Rate) and on the y-axis TPR (True Positive Rate) lies. The formula of this curve measurement is given below :

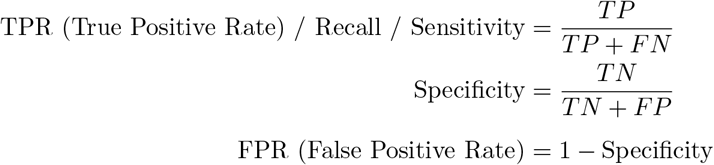

In Figure 4, the AUC-ROC curves illustrate the performance of various classifiers. Each subfigure corresponds to a distinct algorithm, namely Logistic Regression, K-Nearest Neighbor, Support Vector Machine, Decision Tree Classifier, and Random Forest Classifier.

**Figure 4:**
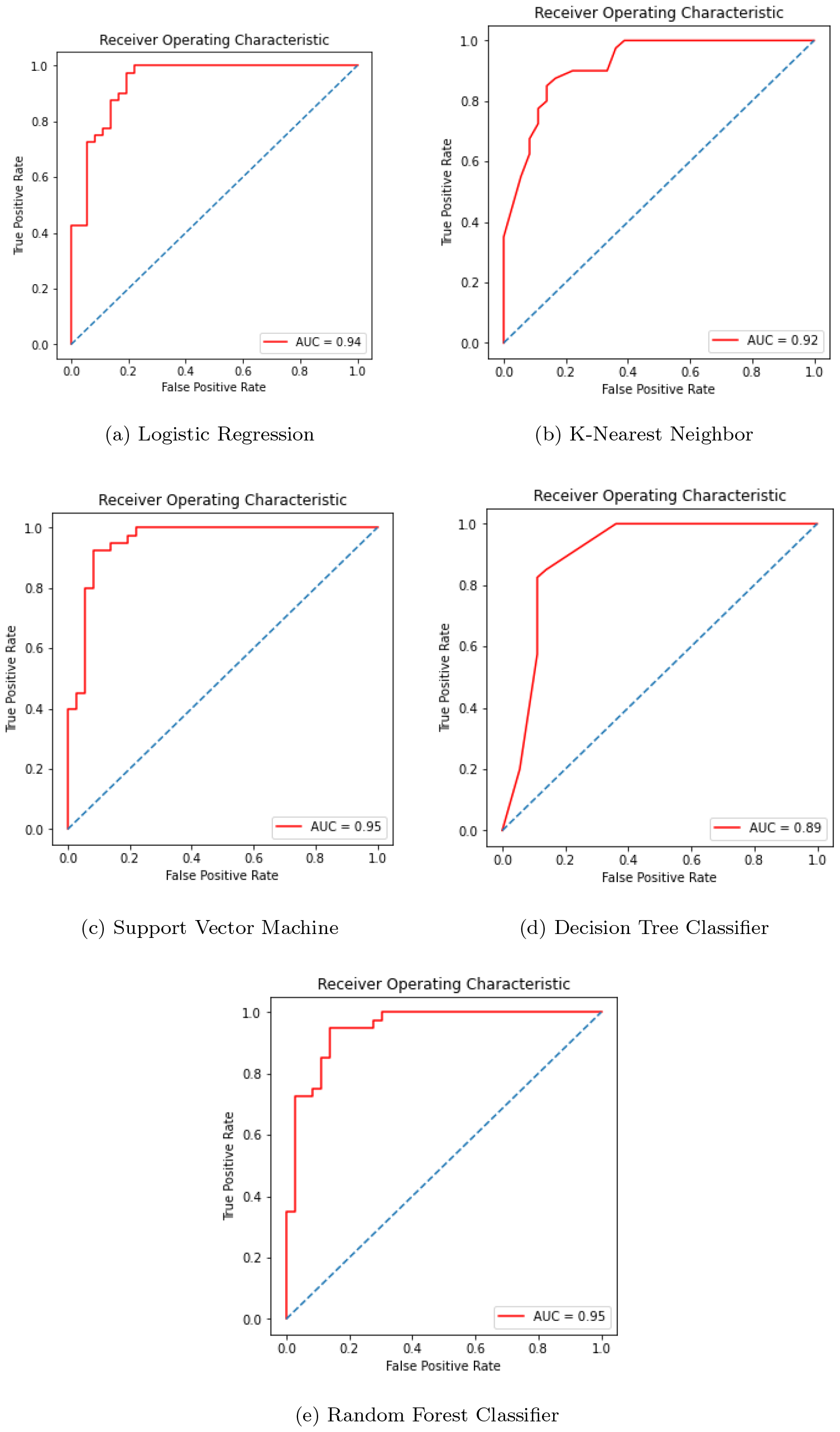
AUC-ROC curves for different classifiers

The AUC values for each classifier are as follows: Logistic Regression (AUC = 0.94), K-Nearest Neighbor (AUC = 0.92), Support Vector Machine (AUC = 0.95), Decision Tree Classifier (AUC = 0.89), and Random Forest Classifier (AUC = 0.95). These AUC values serve as metrics for evaluating the classifiers’ ability to distinguish between positive and negative instances. Higher AUC values generally indicate better discriminatory power, and notably, all the presented classifiers demonstrate very good performance in this regard.

## 7. Analysis and Comparison

Our experiment shows that various algorithms performed better contingent on the circumstance if cross-validation and feature choice are utilized. Each algorithm has its characteristic ability to out-perform others relying on the circumstance. For instance, Random Forest performs much better with an enormous number of datasets than when information is little, while Support Vector Machine performs better with fewer datasets. In the event of a decision tree, missing values play a significant role. Even after ascribing, it can’t give the outcome which it can with an ideal dataset.

We have seen from the model performance tables that after doing hyperparameter tuning, our models perform really well, and also the over-fitting problems that were present before hyperparameter tuning are resolved. Another reason for extraordinary performance is feature selection.

From Tables 14 and 15, we can see that surprisingly the logistic regression works much better than others; it gave 93.44% accuracy, and after that, the Support Vector Machine gave 91.80% accuracy. Other models also work very fine for this dataset. Our models performed better than many other existing works in this field.

**Table 14:** Models Performance without hyperparameter tuning.

| Models | Train Accuracy(%) | Test Accuracy(%) |
| --- | --- | --- |
| Logistic Regression | 87.26 | 81.32 |
| K-Nearest Neighbors (KNN) | 76.89 | 65.93 |
| Support Vector Machine (SVM) | 100.00 | 54.95 |
| Decision Tree Classifier | 100.00 | 73.63 |
| Random Forest Classifier | 100.00 | 81.32 |

**Table 15:** Models Performance with hyperparameter tuning.

| Models | Train Accuracy(%) | Test Accuracy(%) |
| --- | --- | --- |
| Logistic Regression | 87.78 | 93.44 |
| K-Nearest Neighbors (KNN) | 82.64 | 86.89 |
| Support Vector Machine (SVM) | 85.54 | 91.80 |
| Decision Tree Classifier | 83.88 | 90.16 |
| Random Forest Classifier | 91.32 | 88.52 |

## 8. Comparison with Related Works

In this section, we discuss the comparison of our work with some existing related papers. The comparative analysis in Table 16 highlights the performance of various heart disease prediction models proposed in the literature. The existing works did not employ the five different algorithms that we utilized in our study. The comparison reveals that our models achieved better performance compared to the existing works.

**Table 16:** Comparison of Heart Disease Prediction Models.

| Author | LR (%) | KNN (%) | SVM (%) | DT (%) | RF (%) |
| --- | --- | --- | --- | --- | --- |
| B. VenkataLakshmi et al. [6] | - | - | - | 84.013 | 85.034 |
| Jaymin Patel et al. [13] | - | - | - | 56.76 | - |
| K. Gomath et al. [11] | - | - | - | - | 79.90 |
| Zeinab Arabasadi et al. [2] | - | - | - | - | 93.85 |
| Sarangam Kodati et al. [12] | - | - | - | 84.81 | - |
| Our Proposal | 93.44 | 86.89 | 91.80 | 90.16 | 88.52 |

Notably, our proposed model outperforms the others, achieving an accuracy of 93.44%, surpassing the accuracies reported by previous studies. B. VenkataLakshmi et al. [6] and Sarangam Kodati et al. [12] utilized Decision Tree and Random Forest, achieving accuracies of 84.013% and 84.81%, respectively. Zeinab Arabasadi et al. [2] achieved a remarkable accuracy of 93.85% by employing a hybrid model combining Artificial Neural Network and genetic algorithm. These comparisons underscore the effectiveness of our proposed machine-learning model in the context of heart disease prediction.

Our proposed model demonstrates superior performance compared to the referenced models, achieving an accuracy of 93.44%. This outperformance can be attributed to the careful selection and combination of machine learning algorithms. Notably, the Logistic Regression model within our proposal attains a high accuracy of 93.44%, outshining other models. Logistic Regression is known for its efficiency in binary classification tasks, such as predicting the presence or absence of heart disease. The ensemble methods, specifically Random Forest, also contribute to the overall robustness and accuracy of our model by leveraging the strength of multiple decision trees. The combined strength of these models, along with hyperparameter tuning and effective data preprocessing, results in a predictive model that excels in the task of heart disease prediction.

## 9. Conclusion

In this research, we conducted a comprehensive comparison of various machine learning algorithms to predict the presence of cardiovascular heart disease. The primary objective of this study was to assess the accuracy of our machine learning models while also scrutinizing their individual performance characteristics, seeking insights into the variability across different algorithms.

We utilized the Cleveland dataset, sourced from the UCI machine learning repository, comprising 14 attributes and 303 instances. To ensure robust accuracy assessment, we employed 10-fold cross-validation. In our analysis, we considered 13 attributes, segregating those with categorical values, and judiciously removed attributes with minimal importance. Remarkably, after the rigorous implementation of all the algorithms, coupled with hyperparameter tuning, our models demonstrated impressive performance. Notably, ‘Logistic Regression’ emerged as the top-performing model, achieving a remarkable accuracy rate of 93.44%. Following closely, ‘Support Vector Machine’ exhibited strong predictive capabilities, with an accuracy rate of 91.80%.

While our study has yielded valuable insights, it’s important to note that the dataset’s limited size may have slightly constrained the models’ performance. Access to a larger dataset could potentially lead to even more robust results. Nevertheless, the Cleveland dataset remains a widely recognized and reliable resource for research of this nature.

In conclusion, our research underscores the potential of machine learning in the early detection of cardiovascular heart disease. ‘Logistic Regression’ and ‘Support Vector Machine’ have emerged as promising tools for accurate risk assessment. As we continue to refine and expand our methodologies, this research contributes to the broader goal of optimizing healthcare practices and improving patient outcomes.

Looking forward, future work in this domain may involve the exploration of more extensive and diverse datasets to enhance model generalization and robustness. Additionally, fine-tuning and experimenting with advanced machine learning algorithms and ensemble methods could yield even more accurate predictions. Furthermore, the integration of real-time patient data and continuous monitoring through wearable devices holds promise for proactive cardiovascular health management.

## 9.1. Acknowledgement

This research is supported by the Jagannath University Research Grant, Dhaka, Bangladesh.

